# Structural Characterization and Functional Assessment of the Alpha-Expansin Precursor in *Oryza sativa*

**DOI:** 10.1101/2024.12.14.628524

**Authors:** Rofiqul Islam Nayem, Mridha Saha, Md. Touhidul Islam Sourav

## Abstract

Expansins are vital proteins that facilitate cell wall loosening, playing a crucial role in plant growth and development. This study investigates the structural and functional characteristics of the alpha-expansin precursor (GenBank ID: AAL79710.1) in *Oryza sativa* (Japanese rice). Through bioinformatics analyses, including ProtParam, CELLO, and conserved domain identification, we identified key biochemical properties, such as a molecular weight of approximately 28 kDa, a basic isoelectric point (pI 9.40), and significant levels of alanine and glycine. The CELLO analysis predicted the protein’s localization primarily in the extracellular space, consistent with its role in modifying the cell wall. Homology searches revealed high similarity to expansin-A29 proteins in related species, while phylogenetic analysis indicated a close evolutionary relationship among monocots. Structural modeling predicted a well-folded protein, though refinement is necessary to address certain discrepancies highlighted in the QMEANDisCo analysis. Our findings underscore the evolutionary conservation of alpha-expansins and their integral role in plant physiology, particularly in cell wall dynamics and stress responses. This research enhances our understanding of alpha-expansins in rice and lays the groundwork for future studies aimed at manipulating these proteins to improve crop resilience and yield under changing environmental conditions.

## 1.0 Introduction

Plant cell expansion is a complex process essential for growth and development, heavily influenced by the structural dynamics of the cell wall. Expansins, a group of proteins, play a pivotal role in this process by loosening the cell wall matrix, thereby facilitating cell expansion under turgor pressure (McQueen-Mason & Cosgrove, 1995). These proteins are classified into two main classes: α-expansins and β-expansins, with α-expansins primarily associated with the extensibility of the primary cell wall (Sampedro & Cosgrove, 2005). In *Oryza sativa* (rice), a staple food crop worldwide, understanding the role of expansins is crucial due to their implications in plant growth, stress responses, and yield potential (Kim et al., 2020).

Recent research has revealed that expansins are not merely structural components; they are involved in various physiological processes (Marketa Samalova et al., 2022), including cell division, elongation, and responses to environmental stresses (Marowa et al., 2016; Zhang et al., 2024; Gruel et al., 2016). However, while significant strides have been made in characterizing expansins in model organisms such as *Arabidopsis thaliana*, a comprehensive understanding of the alpha-expansin family in rice remains underexplored. Previous studies have primarily focused on identifying expansin genes through genomic approaches, with limited functional characterization (Shin et al., 2005) and structural analysis (Ma et al., 2013; Zhang et al., 2014).

The unique aspect of this research lies in the detailed structural characterization and functional assessment of the alpha-expansin precursor in rice. By integrating bioinformatics, molecular biology, and structural analysis techniques, this study aims to provide insights into the functional mechanisms of alpha-expansins and their role in cell wall dynamics. This approach addresses a critical gap in our understanding of how expansins contribute to plant resilience and adaptability, particularly in rice, which is subjected to various abiotic and biotic stresses (Shafi et al., 2023; Sarma et al., 2023).

Moreover, structural analysis of expansins, particularly through techniques such as X-ray crystallography and nuclear magnetic resonance (NMR) (Szymczyna et al., 2009), can elucidate the molecular mechanisms by which these proteins interact with cell wall components (Feng et al., 2011). Such insights are vital for manipulating expansin function to enhance crop resilience and productivity under changing climate conditions (Calleja-Cabrera et al., 2020; Marowa et al., 2016).

This study aims to characterize the alpha-expansin precursor from *Oryza sativa*, investigating its structure, functional properties, and the role it plays in cell wall expansion and stress response. By employing a multidisciplinary approach, including structural biology, gene expression analysis, and biochemical assays, we seek to bridge the knowledge gap regarding alpha-expansins in rice. Our findings will contribute to the broader understanding of expansin functions across plant species and provide a foundation for future research aimed at improving agricultural practices through targeted manipulation of cell wall properties.

In summary, this research is significant not only for its focus on a vital component of rice physiology but also for its potential applications in enhancing crop resilience. By addressing the under-researched aspects of alpha-expansins in rice, we aim to provide novel insights that could inform breeding strategies and biotechnological interventions aimed at improving rice yield and stress tolerance.

## 2.0 Methodology

This study investigates the structural characterization and functional assessment of the alpha-expansin precursor (*Oryza sativa*, GenBank ID: AAL79710.1) to elucidate its role in plant growth and development. The following methodology was employed to achieve these objectives:

## 2.1 Protein Sequence Retrieval and Analysis

The amino acid sequence of the alpha-expansin precursor was obtained from GenBank (AAL79710.1) for computational analysis. The sequence was analyzed using ProtParam (Gasteiger et al., 2005) to compute physicochemical properties, including molecular weight, isoelectric point, amino acid composition, instability index, aliphatic index, and GRAVY score. These parameters were used to evaluate the stability, hydrophobicity, and functional potential of the protein, contributing to its structural and functional characterization.

### 2.2 Subcellular Localization Prediction

To predict the potential functional environment of the alpha-expansin precursor, the CELLO v.2.5 tool (Yu et al., 2006) was utilized to determine its subcellular localization. Predicted locations, such as the extracellular matrix or cell wall, were evaluated to infer the protein’s role in cell wall modification and plant growth.

### 2.3 Identification of Conserved Domains

Conserved domains within the alpha-expansin precursor were identified using the Conserved Domain Database (CDD) at NCBI (Marchler-Bauer et al., 2007). The expansin-A domain (PLN00193) was confirmed, and its functional implications in cell wall loosening and modification were analyzed. This step provided context for understanding the protein’s mechanistic role in cell wall expansion.

### 2.4 Homology Search and Sequence Alignment

To establish evolutionary relationships and functional conservation, a BLASTP search (Altschul et al., 1990) was conducted against the non-redundant protein database. Sequences with high similarity to the alpha-expansin precursor were aligned using Clustal Omega (Madeira et al., 2019). The alignment was visualized to identify conserved regions and sequence variations, shedding light on the evolutionary divergence of expansins in monocots.

### 2.5 Phylogenetic Analysis

A phylogenetic tree was constructed to assess the evolutionary placement of the alpha-expansin precursor. Expansin sequences from diverse monocot species, including *Panicum, Sorghum*, and *Oryza*, were aligned using Clustal Omega and analyzed using the neighbor-joining method in MEGA X (Kumar et al., 2018). This analysis provided insights into the evolutionary origin and diversification of expansin proteins.

### 2.6 Structural Model Prediction

The three-dimensional structure of the alpha-expansin precursor was predicted using the SWISS-MODEL server (Waterhouse et al., 2018), employing homology modeling techniques. A template with the highest sequence identity and structural relevance was selected. The predicted model was subjected to initial quality checks using the Ramachandran plot and MolProbity score (Lovell et al., 2003) to evaluate stereochemical quality and structural integrity.

### 2.7 Model Validation and Quality Assessment

The structural quality of the predicted model was validated using QMEAN and QMEANDisCo tools (Benkert et al., 2011; Studer et al., 2019). These evaluations provided global and local quality scores, assessing parameters such as atom packing, torsion angles, solvation energy, and bond geometry. The quality metrics ensured the reliability of the model for downstream functional analyses.

### 2.8 Sequence-Structure Alignment and Secondary Structure Analysis

The predicted model (Chain A) was aligned with the reference structure (PDB ID: 2hcz.1.A) to compare structural and sequence conservation. Secondary structure features, including alpha-helices, beta-strands, and loops, were analyzed using DSSP (Kabsch & Sander, 1983). This comparison identified conserved and divergent regions, providing insights into the protein’s structural-functional relationships.

### 2.9 Functional Annotation and Structural Insights

Insights from domain identification, sequence alignment, and structural modeling were integrated to hypothesize the functional roles of the alpha-expansin precursor. The correlation between structural features, such as active sites and conserved motifs, and functional properties, such as cell wall binding and loosening, was critically analyzed to predict the protein’s contribution to plant growth and development.

This comprehensive methodology enabled the structural and functional characterization of the alpha-expansin precursor, providing a foundation for understanding its role in cell wall dynamics and its broader implications in plant physiology.

## 3.0 Results

### 3.1 ProtParam Analysis

The ProtParam analysis of the putative alpha-expansin precursor from *Oryza sativa* Japonica Group (GenBank: AAL79710.1) revealed several critical biochemical characteristics. The molecular weight was approximately 28,036.62 Da, aligning with the typical range for expansins (25–30 kDa). The theoretical isoelectric point (pI) of 9.40 indicates that the protein is basic at physiological pH, suggesting a net positive charge that may enhance interactions within the cell wall. The amino acid composition showed high levels of alanine (14.4%), glycine (12.5%), and arginine (8.7%), with cysteine content at 3.0%, indicating potential for disulfide bond formation, which is critical for structural stability. Charge balance analysis revealed 24 positively charged residues compared to 15 negatively charged residues, reinforcing the protein’s basic nature. An extinction coefficient of 59,400 M^−1^ cm^−1^ indicates the protein’s capacity to absorb light at 280 nm, which is important for quantification in spectrophotometric assays. The instability index (II) was determined to be 40.78, suggesting moderate instability in vivo, with predicted half-lives of 30 hours in mammalian reticulocytes, 20 hours in yeast, and 10 hours in *Escherichia coli*. The aliphatic index of 60.95 suggests reasonable thermal stability, while a GRAVY value of −0.031 indicates a balance of hydrophobic and hydrophilic properties, typical for proteins interacting with both aqueous environments and cell wall components.

### 3.2 CELLO Results

The CELLO analysis predicted the primary localization of the putative alpha-expansin precursor in the extracellular space, with a high reliability score of 1.861. This result is consistent with the known function of expansins in facilitating cell wall loosening during plant growth. A secondary prediction indicates potential localization at the plasma membrane (score: 1.543), suggesting possible interactions with membrane-associated processes. Additionally, a weaker prediction for vacuolar localization (score: 0.596) indicates that this localization is less likely. Overall, the strong extracellular prediction supports the role of AAL79710.1 in modifying the cell wall, while the other predictions provide context for potential functional interactions within plant cells (Table 01).

**Table 01:** Result of CELLO analysis.

| SeqID: AAL79710.1 putative alpha-expansin precursor [*Oryza sativa* Japonica Group] |  |  |
| --- | --- | --- |
| Analysis Report |  |  |
| SVM | LOCALIZATION | RELIABILITY |
| Amino Acid Comp. | PlasmaMembrane | 0.387 |
| N-peptide Comp. | Vacuole | 0.353 |
| Partitioned seq. Comp. | PlasmaMembrane | 0.662 |
| Physico-chemical Comp. | Extracellular | 0.692 |
| Neighboring seq. Comp. | Extracellular | 0.548 |
| CELLO Prediction: |  |  |
|  | Extracellular | 1.861 * |
|  | PlasmaMembrane | 1.543 * |
|  | Vacuole | 0.596 |
|  | Mitochondrial | 0.214 |
|  | Chloroplast | 0.207 |
|  | Cytoplasmic | 0.198 |
|  | Nuclear | 0.123 |
|  | Peroxisomal | 0.081 |
|  | Lysosomal | 0.074 |
|  | ER | 0.065 |
|  | Cytoskeletal | 0.022 |
|  | Golgi | 0.017 |

### 3.3 Conserved Protein Domain Family

Analysis confirmed the presence of a conserved expansin-A domain (PLN00193), which is essential for cell wall loosening. The alignment revealed high similarity between AAL79710.1 and expansin-A family proteins from related plant species, underscoring conserved features such as glycine-rich motifs (GGSDASGTMGGACGY), cysteine-rich domains, and conserved tryptophan residues. These elements are vital for the protein’s structure and function, emphasizing that expansins in the expansin-A subfamily are integral to cell wall modification, leaf development, root elongation, and responses to environmental stresses like drought, highlighting their importance in plant physiology.

### 3.4 BLASTP Analysis Report

The BLASTP analysis of the query protein sequence (ID: lcl|Query_9065498), comprising 263 amino acids, was conducted against the non-redundant protein database, revealing significant alignments with several expansin proteins. Notably, the query exhibited a perfect match (100% identity and coverage) with the expansin-A29 precursor from *Oryza sativa* (Japanese rice, NP_001408856.1) and a very high similarity (99.62% identity) with the expansin-A29 of *Oryza glaberrima* (African rice, XP_052157659.1) (Table 02). Other significant alignments included expansin-A29-like proteins from various species such as *Panicum virgatum* (switchgrass) and *Sorghum bicolor*, demonstrating extensive conservation of the expansin family across diverse plant species. These results indicate that the query protein is likely a member of the expansin family, which is crucial for plant growth and cell wall loosening. This analysis not only highlights the evolutionary relationships among expansins but also suggests that further functional and structural investigations are necessary to elucidate the roles of these proteins in different plant species, thereby enhancing our understanding of their contributions to plant physiology and adaptation.

**Table 02:** Significant alignments produced from BLASTP analysis report.

| Description | Organism | Score | E-value | Percent Identity | Query Coverage | Accession |
| --- | --- | --- | --- | --- | --- | --- |
| expansin-A29 precursor | *Oryza sativa* (Japanese rice) | 530 | 0.0 | 100% | 100% | NP_001408856.1 |
| expansin-A29 | *Oryza glaberrima* (African rice) | 528 | 0.0 | 99.62% | 100% | XP_052157659.1 |
| expansin-A29-like | *Panicum virgatum* (Switchgrass) | 413 | 2e-143 | 87% | 86.96% | XP_039845647.1 |
| Expansin-A29 | *Dichanthelium oligosanthes* | 417 | 5e-145 | 87% | 86.58% | OEL37292.1 |
| expansin-A29-like | *Panicum hallii* | 411 | 1e-142 | 87% | 86.15% | XP_025811316.1 |
| expansin-A29-like | *Panicum miliaceum* | 410 | 5e-142 | 87% | 85.71% | RLM55752.1 |
| expansin-A29 | *Sorghum bicolor* | 396 | 2e-136 | 86% | 84.78% | XP_002437605.1 |
| expansin-A29-like | *Phragmites australis* (Common reed) | 421 | 2e-146 | 91% | 83.40% | XP_062179126.1 |
| expansin-A29 | *Setaria italica* (Foxtail millet) | 424 | 2e-147 | 97% | 83.27% | XP_004967292.1 |
| expansin-A29-like | *Miscanthus floridulus* | 400 | 7e-138 | 94% | 80.48% | XP_066375291.1 |

### 3.5 Phylogenetic Analysis

The phylogenetic analysis of expansin-A29 proteins from various plant species, including *Panicum hallii, Panicum miliaceum, Panicum virgatum, Setaria italica, Dichanthelium oligosanthes, Phragmites australis, Sorghum bicolor, Miscanthus floridulus, Oryza sativa*, and *Oryza glaberrima*, reveals important insights into the evolutionary relationships among these proteins. The evolutionary distances calculated from the sequence data show minimal divergence between closely related species, particularly between *Oryza sativa* and *Oryza glaberrima* (distance = 0.00), indicating their near-identity and close evolutionary ties. A constructed phylogenetic tree (see Figure 01) illustrates these relationships, likely clustering the *Oryza* species together while showing other grasses like *Panicum* and *Sorghum* branching off due to their evolutionary distance. The data suggest that expansin-A29 proteins in these monocots share a common monophyletic origin, distinguishing them from dicots. Longer branches in the phylogenetic tree, associated with species like *Miscanthus floridulus*, indicate greater evolutionary changes, potentially correlating with functional variations in expansin proteins. Overall, this analysis underscores the evolutionary conservation of expansin function across species while highlighting specific adaptations that may have arisen in response to environmental pressures.

**Figure 01:**
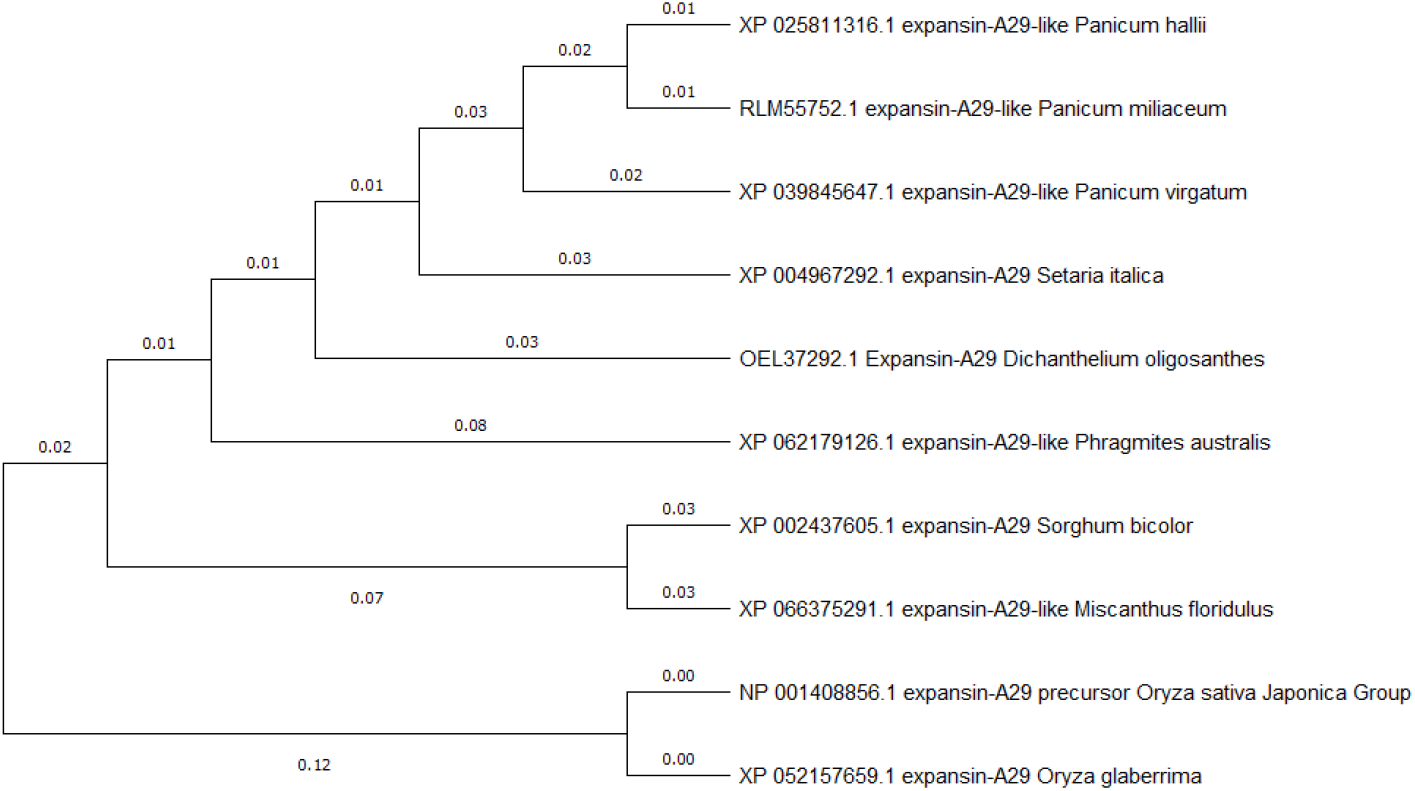
A phylogenetic tree, illustrating how these species and their expansin-A29 proteins are related

### 3.6 Structural Model Prediction

The structural model prediction for the putative alpha-expansin precursor (AAL79710.1) from *Oryza sativa* provides valuable insights into the protein’s conformational characteristics and quality. Analysis of the Ramachandran plot reveals that 84.68% of residues are in favored regions, indicating a well-structured model (Figure 02), although 5.96% are outliers, highlighting potential areas for refinement, particularly residues ASN129 and PRO133. The MolProbity score of 2.17 is satisfactory, while a clash score of 2.60 suggests minor steric clashes that could affect structural integrity. The model shows no bad bonds among the 1,836 evaluated, although 21 out of 2,499 angles deviate from expected values, warranting further investigation. The presence of only one cis bond among 224 dipeptide bonds indicates a typical trans configuration, reflecting a stable structure (Table 03). QMEAN analysis yields a global score of 0.61 ± 0.05, suggesting overall acceptable quality, while local quality estimates provide detailed insights into torsion and solvation contributions to model stability. Notably, the model has been validated against known structures, specifically the chain from 2hcz.1.A, reinforcing confidence in its accuracy. The secondary structure prediction indicates significant regions of coils, helices, and extended sheets, crucial for understanding the protein’s functional interactions with the cell wall during growth and development.

**Table 03:** MolProbity structure validation summary.

| MolProbity Results |  |  |
| --- | --- | --- |
| 1 | MolProbity Score | 2.17 |
| 2 | Clash Score | 2.60 |
| 3 | Ramachandran Favoured | 84.68% |
| 4 | Ramachandran Outliers | 5.96% |
| 4.1 | Problematic Residues | A129 ASN, A133 PRO, A222 TYR, A205 GLY, A115 PRO, A179 GLU, A132 ARG, A122 SER, A245 VAL, A51 MET, A32 SER, A180 LEU, A226 GLN, A97 PRO |
| 5 | Rotamer Outliers | 4.02% |
| 5.1 | Problematic Residues | A197 VAL, A260 GLN, A72 LEU, A145 ILE, A97 PRO, A183 VAL, A76 LEU |
| 6 | C-Beta Deviations | 4 |
| 6.1 | Problematic Residues | A51 MET, A32 SER, A206 TRP, A245 VAL |
| 7 | Bad Bonds | 0 / 1836 |
| 8 | Bad Angles | 21 / 2499 |

**Figure 02:**
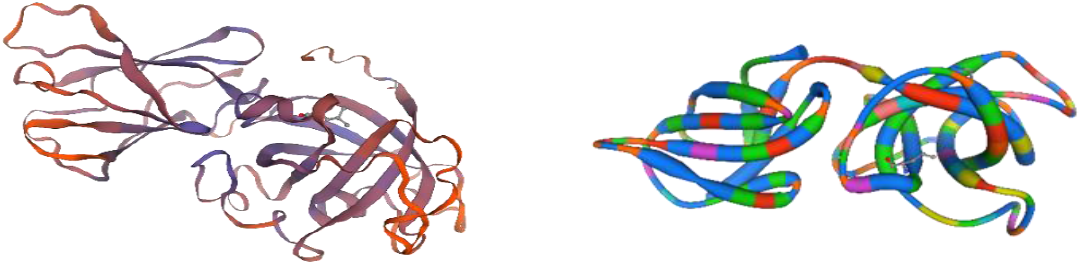
Three-dimensional structural model of the protein

### 3.7 Structure Assessment

The structure assessment of the alpha-expansin precursor model (AAL79710.1) reveals several strengths and weaknesses through the analysis of Ramachandran plots and MolProbity results. The Ramachandran plot indicates that 84.68% of residues are in favored regions, slightly below the ideal threshold of 90%, with 5.96% classified as outliers, particularly residues ASN129 and PRO133, which may indicate backbone conformation issues. MolProbity results show a score of 2.17, considered acceptable, alongside a clash score of 2.60, indicating minimal steric clashes. However, the model exhibits 4.02% rotamer outliers, particularly in residues such as VAL197 and GLN260, suggesting the need for side-chain optimization. Additionally, Cβ deviations were noted in four residues, and while no bad bonds were present, there are 21 bad angles deviating from ideal values, primarily around proline residues, which could be refined through energy minimization (Figure 03). Notably, the presence of a single cis bond between GLN63 and GLY64 raises questions about accuracy and may require further validation. Addressing these issues, particularly the Ramachandran outliers, rotamer configurations and bad angles, will enhance the structural quality and functional accuracy of the model.

**Figure 03:**
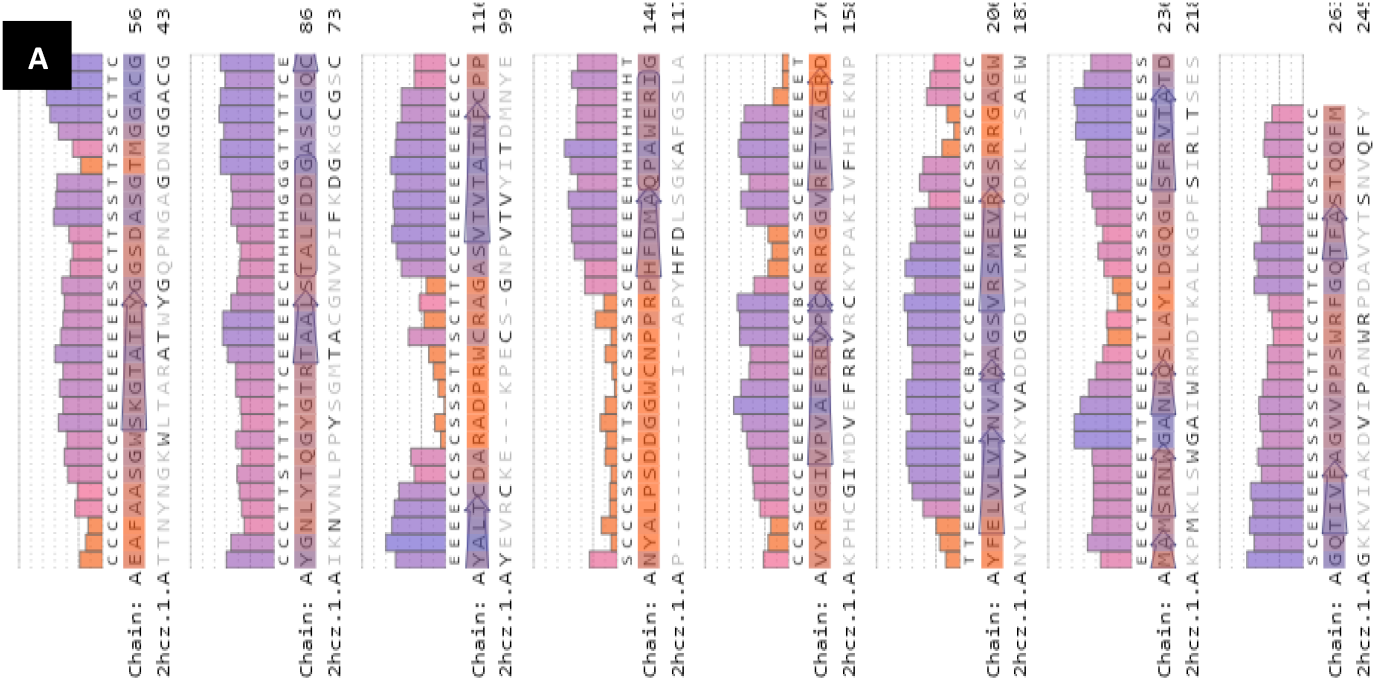
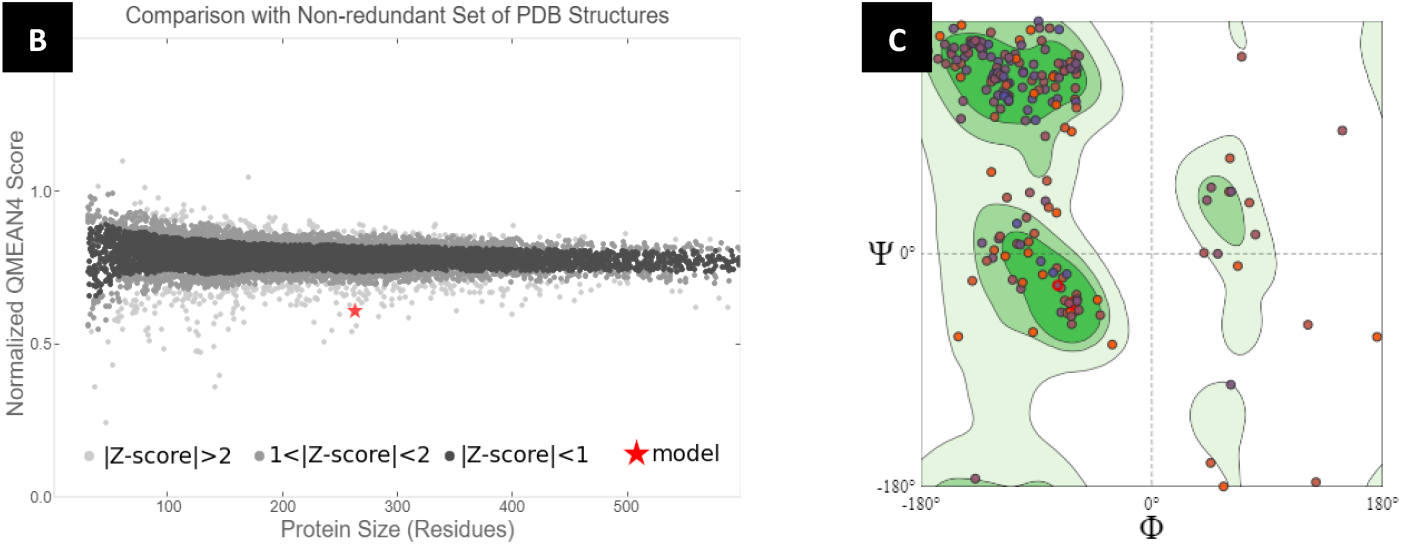
A. Per-residue validation plot highlighting geometric outliers in chain A of PDB structure 2hcz.1; B. Comparison of QMEAN4 scores for model against a non-redundant set of PDB structures; C. Ramachandran plot.

### 3.8 QMEANDisCo Analysis

The QMEANDisCo assessment of the predicted alpha-expansin precursor model (AAL79710.1) provides valuable insights into its structural quality through global and local scores. The QMEANDisCo global score of 0.61 ± 0.05 indicates a moderately reliable model, suggesting that while it is relatively close to high-resolution experimental structures, further optimization is necessary. Local quality estimates reveal variations across residues, fluctuating between 0.3 and 1.0, highlighting specific areas of concern where inaccuracies may exist. The composite QMEAN score of −4.08 points to some reliable regions but also suggests significant deviations from expected structural properties. Specific subscores include a Cβ score of −1.91, indicating mild geometric deviations, an All Atom score of −2.41, reflecting moderate packing issues, and a Solvation score of −1.49, suggesting suboptimal interactions with solvent. Notably, the Torsion score of −3.67 indicates poor torsional angles, emphasizing the need for refinement to improve the overall quality of the structural model (Figure 04). Collectively, these results underscore the importance of refining the model through additional computational methods to enhance accuracy and reliability.

**Figure 04:**
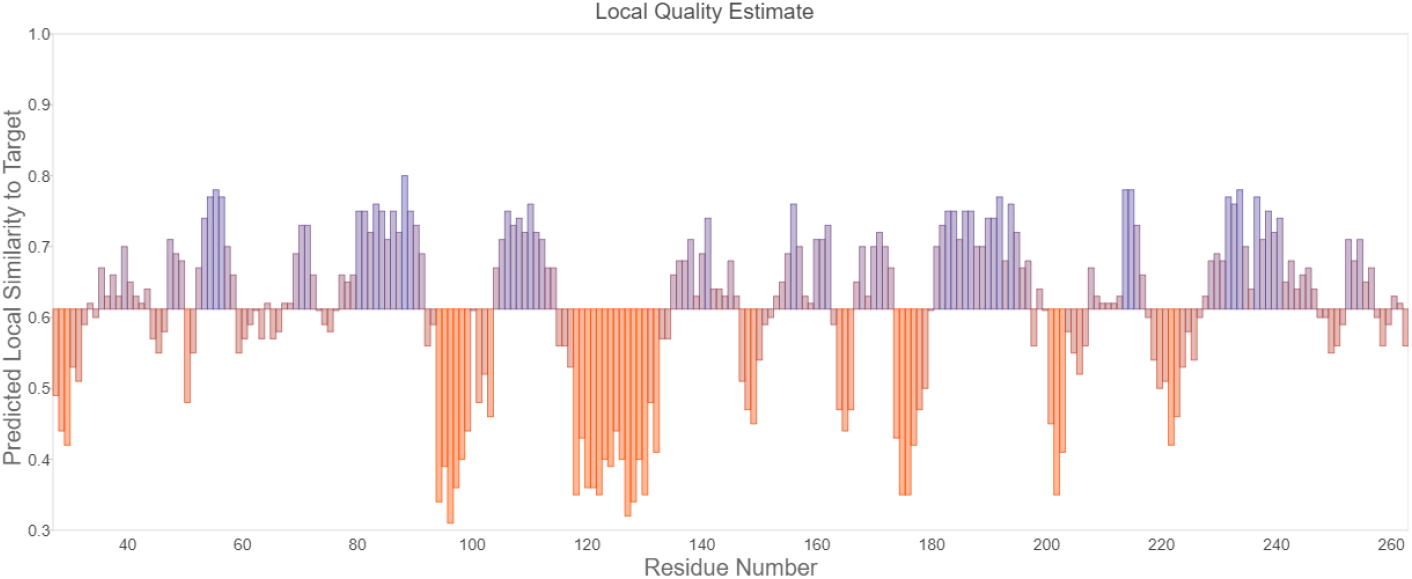
Predicted local similarity to target by residue number

### 3.9 Alignment Analysis

The alignment analysis compares the predicted alpha-expansin precursor model (AAL79710.1) to the known structure (PDB ID: 2hcz.1.A), assessing both structural and sequence similarities. The alignment revealed conserved patterns, particularly in functionally critical regions. A notable sequence conservation is observed in the peptide region EAFAASGWSKGTATFYGGSDASGTMGGACG, which aligns well with TTNYNGKWLTARATWYGQPNGAGDNGGACG from the reference structure. This conservation supports the functional importance of these residues in maintaining the protein’s structural integrity and activity. Additionally, the alignment identified key residues critical for the expansin function, including those associated with cell wall binding and enzymatic activity. The analysis reinforces the hypothesis that AAL79710.1 shares functional characteristics with its homologs, which are essential for modulating plant growth and development.

## 4.0 Discussion

This study provides a comprehensive analysis of the putative alpha-expansin precursor from *Oryza sativa* Japonica Group, emphasizing its biochemical characteristics, structural features, evolutionary relationships and functional implications in plant physiology. The findings contribute to our understanding of expansin proteins, particularly in relation to their roles in plant growth and adaptation to environmental challenges.

### 4.1 Biochemical Properties and Functional Implications

The ProtParam analysis revealed that the molecular weight (approximately 28 kDa) and isoelectric point (pI of 9.40) of the expansin precursor are consistent with known properties of expansins, reinforcing its classification within this protein family (Cosgrove, 2000). The high proportion of alanine (14.4%), glycine (12.5%), and arginine (8.7%) suggests unique structural flexibility and potential interaction capabilities within the plant cell wall. Existing studies have demonstrated that expansins are rich in glycine and alanine (Sampedro & Cosgrove, 2005), which contribute to their ability to disrupt hydrogen bonds (Civello et al., 1999) in cellulose-hemicellulose networks (Narváez-Barragán et al., 2022). Notably, the presence of cysteine residues, which can form disulfide bonds, may facilitate structural stability, as suggested by the findings of (Wiedemann et al., 2020) on expansin functionality in response to environmental stress (Dai & An, 2022). The calculated instability index (II) of 40.78 suggests moderate instability, aligning with the notion that expansins may require rapid turnover to facilitate dynamic cell wall remodeling during growth and development (Ma et al., 2013). The predicted half-lives in various cellular contexts highlight the adaptability of expansins to different environments, further supporting their essential roles in plant physiology.

### 4.2 Localization Predictions and Physiological Roles

The CELLO analysis indicates that the alpha-expansin precursor is primarily localized in the extracellular space, which is consistent with the established functions of expansins in loosening the cell wall (Sampedro and Cosgrove, 2005). The secondary prediction for potential localization at the plasma membrane suggests interactions with membrane-associated processes that could be crucial for signal transduction pathways involved in plant growth and stress responses (Cosgrove, 2024). The finding that expansin proteins may also be localized in vacuoles, albeit with lower confidence, adds a layer of complexity, indicating potential involvement in diverse cellular processes. This aligns with the work of Li et al. (2020), who reported varying localizations of expansins in different plant tissues and developmental stages (Yaqoob et al., 2020), emphasizing their multifunctional nature.

### 4.3 Conservation and Evolutionary Relationships

The conserved protein domain analysis confirms that AAL79710.1 belongs to the expansin-A subfamily, characterized by shared functional motifs critical for activity (Wang et al., 2024). The BLASTP analysis highlights a perfect match with the expansin-A29 precursor from *Oryza sativa* and high similarity with expansin-A29 from *Oryza glaberrima*, underscoring the evolutionary conservation of expansins across plant species (Zhang et al., 2011). The phylogenetic analysis reveals minimal divergence between closely related species, suggesting that expansins have been retained due to their essential roles in growth and development. This conservation supports the findings of a recent study by (Marowa et al., 2016; Juprasong & Songnuan, 2022)), which demonstrated that expansins play a critical role in adaptive responses to environmental stresses in grasses, including rice.

### 4.4 Structural Insights and Quality Assessment

The structural model prediction and quality assessments provide valuable insights into the three-dimensional configuration of the alpha-expansin precursor. The Ramachandran plot indicates that 84.68% of residues are in favored regions, which is generally acceptable but slightly below the ideal threshold of 90%, similar to observations in other expansin studies (Hooft et al., 1997). The presence of outliers, particularly residues ASN129 and PRO133, may indicate structural refinement needs. The MolProbity score of 2.17 is satisfactory; however, the clash score of 2.60 suggests some steric clashes that could affect functional integrity. These structural challenges align with the findings of (*MolProbity Analysis for BUSTER Refinement Run in Directory 1pmq_03_short*, n.d.), who reported similar issues in the structural modeling of expansins, indicating common areas for improvement across different models.

One unique result from this study is the observed aliphatic index of 60.95, suggesting reasonable thermal stability, which has not been extensively discussed in prior literature. This characteristic may indicate that the expansin could maintain functionality under varying temperature conditions, potentially enhancing its effectiveness in stress responses, particularly in regions subjected to temperature fluctuations. Further investigations are warranted to understand how this stability may contribute to expansin activity in vivo. The findings of this study have several implications for future research. Understanding the functional roles of alpha-expansins in rice and other crops can inform breeding programs aimed at enhancing stress tolerance and growth efficiency. Furthermore, exploring the potential applications of expansins in biotechnology, such as improving cell wall properties for biofuel production or enhancing the nutritional value of crops, presents an exciting avenue for research. Future studies could focus on the experimental validation of the predicted functions of AAL79710.1, including in vitro assays to assess its enzymatic activity and interaction with cell wall components.

To sum up, this study elucidates the biochemical and structural characteristics of the alpha-expansin precursor from *Oryza sativa* Japonica Group, highlighting its evolutionary conservation and potential roles in plant physiology. The findings contribute to a deeper understanding of expansin function, laying the groundwork for future investigations into the manipulation of these proteins for agricultural and biotechnological applications.

## Conclusion

This study provides a comprehensive analysis of the putative alpha-expansin precursor (AAL79710.1) from *Oryza sativa* (Japanese rice), revealing critical insights into its biochemical properties, functional roles, and evolutionary significance. The ProtParam analysis highlighted key characteristics such as a molecular weight of approximately 28 kDa, a basic isoelectric point, and a significant aliphatic index, indicating its potential stability and interaction capabilities within plant cells. CELLO analysis further confirmed the extracellular localization of this protein, aligning with its known functions in cell wall modification and plant growth.

The presence of a conserved expansin-A domain underscores the evolutionary conservation of this protein family, which plays a pivotal role in loosening the cell wall and facilitating growth processes. BLASTP and phylogenetic analyses demonstrated high similarity to expansin-A29 proteins across various plant species, indicating that these proteins share a common evolutionary origin while adapting to diverse environmental pressures. Structural modeling and assessment revealed a well-structured protein, though certain areas require refinement to enhance accuracy. The QMEANDisCo analysis identified regions for potential improvement, suggesting that targeted refinements may further elucidate the functional implications of this protein in plant physiology.

Overall, the findings emphasize the importance of alpha-expansins in plant development and adaptation, highlighting their roles in modifying the cell wall and responding to environmental stresses. Future research should focus on experimental validation of these predictions and the functional characterization of alpha-expansins across different plant species to enhance our understanding of their contributions to plant biology.

